# Large scale, simultaneous, chronic neural recordings from multiple brain areas

**DOI:** 10.1101/2023.12.22.572441

**Authors:** Maxwell D Melin, Anup Khanal, Marvin Vasquez, Michael B Ryan, Anne K Churchland, Joao Couto

## Abstract

Understanding how brain activity is related to animal behavior requires measuring multi-area interactions on multiple timescales. However, methods to perform chronic, simultaneous recordings of neural activity from many brain areas are lacking.

Here, we introduce a novel approach for independent chronic probe implantation that enables flexible, simultaneous interrogation of neural activity from many brain regions during head restrained or freely moving behavior. The approach, that we called *indie* (<u>in</u>dependent <u>d</u>ovetail implants for <u>e</u>lectrophysiology), enables repeated retrieval and reimplantation of probes. The chronic implantation approach can be combined with other modalities such as skull clearing for cortex wide access and optogenetics with optic fibers. Using this approach, we implanted 6 probes chronically in one hemisphere of the mouse brain.

The implant is lightweight, allows flexible targeting with different angles, and offers enhanced stability. Our approach broadens the applications of chronic recording while retaining its main advantages over acute recordings (superior stability, longitudinal monitoring of activity and freely moving interrogations) and provides an appealing venue to study processes not accessible by acute methods, such as the neural substrate of learning across multiple areas.

## Introduction

Interactions between brain areas are critical for neural computations that drive a wide range of behaviors. Multi-electrode arrays allow recording from multiple areas simultaneously and, due to recent developments and the integration with CMOS technology, can now be deployed at scale. These advances have enabled cross-area recordings with high yield (e.g. (Allen et al., 2019; Durand et al., 2023; Jun et al., 2017; Steinmetz et al., 2021, 2019)). Nonetheless, major challenges remain for monitoring neural activity chronically (over many days) with multiple probes. Here, we surmount these challenges with a novel approach to implanting and recovering multiple probes for chronic experiments.

Over the last few decades, the number of electrodes deployed to record neural activity has increased from less than a handful to several thousand per probe (Stevenson and Kording, 2011). These advances directly impact the number of neurons that can be recorded simultaneously and have enabled many studies that describe simultaneous population activity in multiple brain areas (Steinmetz et al., 2019; Stringer et al., 2019; de Vries et al., 2020; Wang et al., 2023). However, in most of these studies, the probes are introduced on the day of recording and retracted at the end of the session (commonly referred to as the “acute” recording configuration). This limits the ability to study activity as it changes during the time course of days to months, introduces experimental delays that may compromise behavior performance, and cannot be used in freely-moving behaviors. Existing approaches to record neuronal activity chronically in rodents are either prohibitively expensive or impose constraints on the areas that can be simultaneously measured. For example, one approach is to irreversibly cement probes to the skull (Okun et al., 2016; Krupic et al., 2018; Mimica et al., 2023; Steinmetz et al., 2021) which provides long and stable recordings; however the probes cannot be recovered. This limits the number of probes that experimenters are willing to implant in one subject and makes these experiments only within reach of a select group of labs. Indeed, most labs consider Neuropixels a precious resource and have a sufficiently limited supply that re-using probes is a necessity. An alternative approach to irreversible cementing is to secure the probe(s) to a fixture, usually 3D-printed, that is attached to the skull (Juavinett et al., 2019; Luo et al., 2020; van Daal et al., 2021; Jones, 2023; Bimbard et al., 2023; Horan et al., 2024). There are a wide range of implant strategies; existing fixtures have been used to implant 1-2 probes in mice and 1-3 in rats.

Recoverability is possible and has been reported for many of the designs. However, existing designs are not ideally suited for multi-probe experiments (e.g. probes are sometimes glued to each other in multi-probe implants, limiting implantation geometries and constraining future experiments to fixed configurations). In detail, Juavinett et al. (2019) reports 4 successful explants out of 10 insertions with 2 re-uses. Luo et al. (2020) report 8 successful explants out of 22 insertions with 3 reuses of 2 probes. Steinmetz et al. (2021) reports 4 reuses with 2 probes for periods of 2 weeks. Two recent multi-lab efforts were able to further characterize their implant strategies by performing >60 insertions. Bimbard et al. (2024) reports 55 out of 63 successful explants and 30 re-uses. Horan et al. (2024) reports 127 explants out of 175 insertions with 36 re-uses. It remains unclear, however, whether using probes for many months impacts re-usability or extraction success since these metrics are inconsistently reported across studies. Further, when many probes are used in the same animal or when dealing with a limited supply of probes, implant recoverability becomes critical.

Several additional constraints limit the flexibility and adoption of published designs for chronic, multi-probe fixtures. First, in some cases, probes are attached to the same fixture and lowered together, as a unit, into the brain, as in (van Daal et al., 2021; Bimbard et al., 2023). This approach restricts the areas that can be targeted because it does not allow probes to be driven at independent angles, and imposes a minimum distance between the probes (3mm in van Daal et al. 2021). Weight is the second critical consideration for multiprobe implants: animals can only carry a fraction of their body weight. Implants that weigh more than 15-20% of the animals’ weight are prohibitive. Third, fixtures are often composed of multiple parts that have complicated assembly protocols, e.g. (Jones, 2023; van Daal et al., 2021), making fixture assembly time consuming. Lastly, the duration of the surgery is a critical factor when implanting multiple probes. It is common that surgery times extend 4-6h for 2 probes. However, the duration of surgical procedures is often not reported and likely to vary from user to user. Approaches that allow implanting multiple probes at independent angles are limited and pose constraints on the number of probes or steepness of the angles used because additional shielding is required (Jones, 2023).

We set out to develop a novel fixture implant and surgical strategy that overcomes current limitations. Our design enables studying brain activity on long timescales, provides high targeting flexibility at reduced weight, affords high reuse rates, and permits shortened assembly and surgical times, using only a single mechanical structure. Using this approach, we were able to simultaneously record with 6 probes (24 shanks) from selected targets in one hemisphere of the mouse brain. Recordings are also stable for over 310 days, thus enabling long-term investigation of neural activity from multiple areas at the same time.

## Results

### Novel fixture for multiprobe implants with optimized surgical procedures

Our approach takes advantage of commercially available Neuropixels probes (Jun et al., 2017; Steinmetz et al., 2021). We report results using 384 recording sites per probe; the approach is scalable to the higher channel devices currently under development. Neuropixels probes are currently available with a dovetail attached. This feature has not been exploited by existing chronic implant solutions; we established procedures to take advantage of the dovetail and thus simplify the assembly and probe explant. Importantly, securing Neuropixels via the dovetail allows for fast assembly, easy explant (by sliding the probe out along the dovetail socket), and reuse of probes for acute experiments or chronic experiments requiring different holder dimensions (because probes are not glued to the holder). We engineered a probe fixture (Fig. 1a and Supplementary Fig. 1a,b) that can be 3D printed using stereolithography (SLA) technology on a desktop printer. The use of a desktop printer (e.g. FormLabs, Form 3+) allows individual labs to extend/adapt the design with ease. The dovetail socket in the fixture requires small tolerances that are at the limit of current SLA technology; we provide detailed instructions to reliably manufacture holders (see Methods).

**Figure 1.**
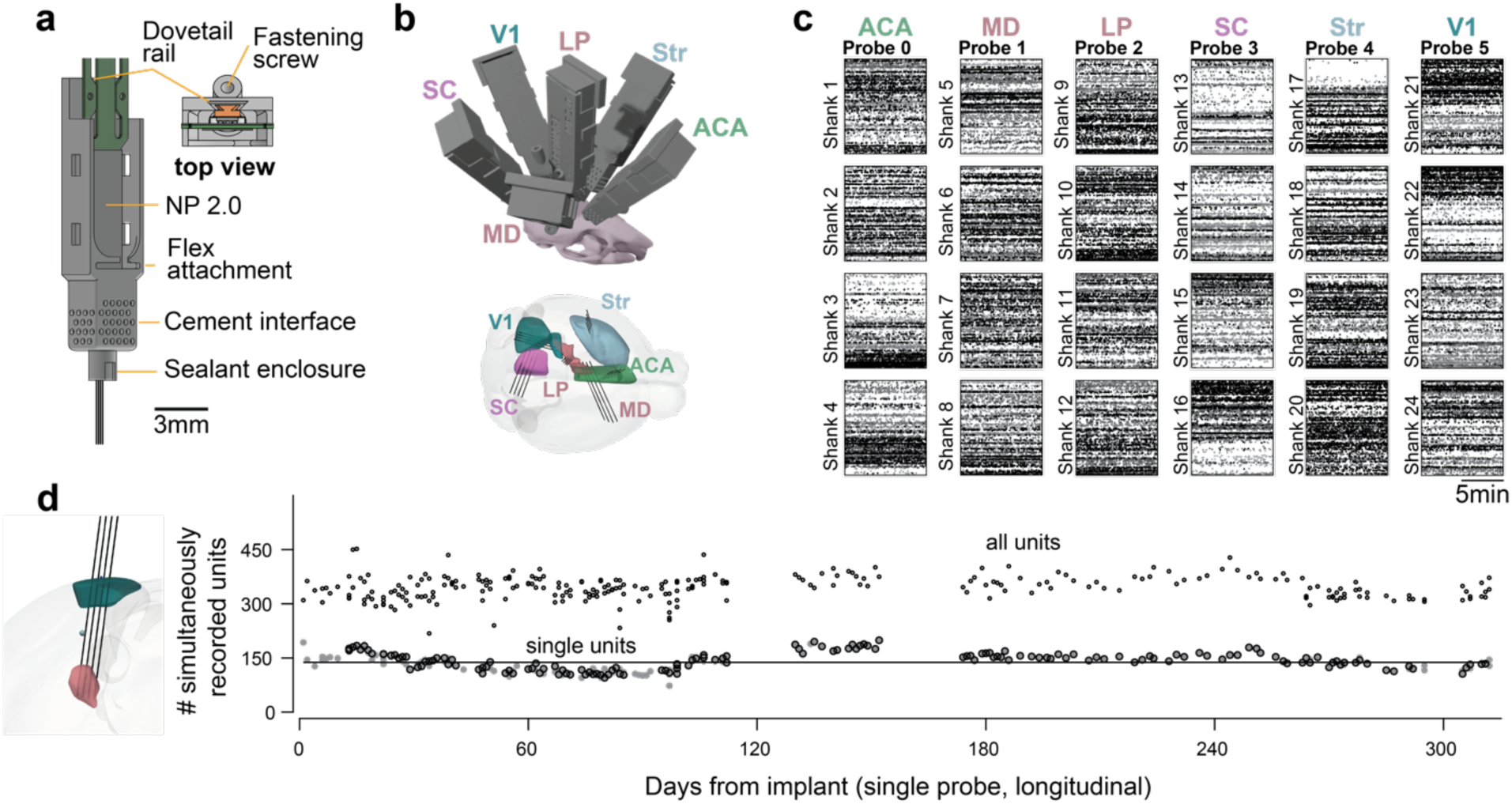
Novel fixture for chronic, large-scale electrophysiology. a) Neuropixels 2.0 probe in fixture and description of details. b) Simultaneous chronic 6 probe implant in the same hemisphere of a mouse targeting specific structures. The weight of the implant is 3.42g including the probes and all fixtures. c) Spike rasters for the 24 shanks implanted in (b). Color indicates spike amplitude (black is high amplitude); abscissa is time, ordinate is depth from the selected recording area in each shank. (d) Left: Planned trajectory of a 4-shank Neuropixels probe targeting cortex (green) and thalamus (pink). Right: Single unit yield is maintained for >300 days. Number of simultaneously recorded units for a mouse implanted with a probe from the day of implant. Line: average. Open circles: all units. Gray filled circles: single units. Dots with black border were recorded during a perceptual decision-making task, while dots without the border were recorded in passive conditions. All circles are recorded from the same probe configuration and represent the number of simultaneously recorded units.

The probe is housed within a fixture that forms the primary structure of the assembly (Fig. 1a). During stereotaxic implantation, the fixture is cemented to the skull, and a covering/cap is attached after implantation. At the end of the experiment (weeks to months later) the probe can be released from the fixture and recovered. Importantly, the fixture consists of a single 3D printed structure that forms the sole mechanical structure of the probe holder – the covering/cap offers protection but does not contribute to stabilizing the electrode. This design choice eliminates structural fastenings between 3D printed parts that are often used in existing approaches and can hinder stability.

Multiple probes can be implanted using individual fixtures (at a weight-cost of 0.57g per probe - including the probe, chronic holder, and cap); this allowed us to implant 6 probes in a single hemisphere of the mouse brain (see Supplementary Fig. 1d for examples of implants). After recovery from surgery, mice carry the implant with ease in the home cage (total weight of 3.42g including 6 probes, fixtures and coverings/caps – excluding 1-2g of cement). While the full, 6-probe configuration may impact behavior in freely moving experiments (more on this below), it is entirely feasible for head-restrained experiments in which the animal need not bear the weight of the headstages during the experiment. Using this approach, we were able to simultaneously record 1206 single units (3880 including multi-unit) across 6 target structures (Fig. 1b, c - 1 mouse). Further, because probes have switchable sites (Jun et al., 2017), we can access cells in structures distributed along the insertion tracts across different recording days (not shown). In additional experiments, we characterized the long-term stability of our implants. Recording from thalamus and primary visual cortex (trajectory shown in Fig. 1d), we quantified the unit yield from our device and found that we can reliably record from similar numbers of single units for over 300 days (Fig. 1e).

In addition to designing a novel fixture, it was essential to optimize the surgical procedures for multi-probe implants (Supplementary Fig. 1c). Chronic surgical procedures are rarely standardized across labs. However certain principles, such as the use of viscous cement to carefully create a well around the implant, leaving the shank open to air, are relatively constant across published work (Bimbard et al., 2023; Horan et al., 2024; Jones, 2023; Juavinett et al., 2019; Luo et al., 2020; van Daal et al., 2021). When implanting a single probe, or multiple probes attached to the same fixture, surgeries are reported to take 4-5 hours (van Daal et al., 2021), which would render implanting multiple probes on independent fixtures impractical and time consuming. Long surgery times also complicate animal recovery and introduce steep adoption curves for novice users. We therefore sought to simplify the procedure by encapsulating the shank(s) in silicone adhesive so cement could be added ad-lib. We chose a low toxicity silicone adhesive (KwikSil, WPI) because it can be directly applied to craniotomies, hardens quickly, and provides electrical insulation. However, KwikSil allows the shank(s) to slide through it without breaking during extraction. To facilitate the application of KwikSil, we include a sealant enclosure (Fig. 1a) in the fixture design that ensures the sealant does not leak close to the base of the probe (where probe motion could lead to breakage). With this design, the sealant offers protection to the probe shank(s), and it can be applied without a microscope. Importantly, because the shanks are completely covered by silicone, cement can then be applied ad-lib to secure the fixture in place. We included holes in the housing that act as interfaces to the cement but keep it away from the probe by surface tension (Fig. 1a). Optimizing the surgical procedures greatly reduced the time required to implant multiple probes. An experienced surgeon can implant 6 probes in roughly 6 hours.

Altogether, by creating a novel fixture and optimizing the surgical procedures we could implant multiple probes targeting many distinct brain areas in a single hemisphere of the mouse brain. Importantly, the novel fixture allows retrieval of the probe for reuse in chronic or acute experiments.

### Implant stability and comparison with a published dataset

We established a recoverable method for implanting electrodes that enables recording neural activity chronically. Ideally, the stability of measurements with our fixture would match that of cemented probes and the unit yield would be on par with acute datasets, where the chances of loss of units due to inflammatory response are minimal.

First, we investigated recording stability by quantifying brain motion in relation to the shank. A major concern in electrophysiological experiments is that motion of the brain in relation to the probe causes drift of the recorded neurons during individual sessions. This is problematic when experiments require isolating spikes from individual neurons, such as when measuring pairwise correlations. In our approach, covering the probe and craniotomy with silicone sealant could in principle reduce the amount of motion, leaving less room for the brain to move in the dorsal-ventral direction and securing the shank(s). We therefore set out to quantify the motion of the brain in relation to the shank(s) by using the recorded voltage signals. We implanted a Neuropixels 2.0 alpha probe with 4 shanks, and recorded from 2 shanks in cortex and 2 shanks in thalamus. The recordings were done in a head restrained configuration, with the mouse allowed to run on a treadmill. We reasoned that a condition where the mouse is locomoting vigorously and the skull is fixed would be more likely to produce large artifacts than in a freely moving configuration. We deployed DREDge (Windolf et al., 2023), a motion registration algorithm for electrophysiological data that has been extensively validated in similar recordings with imposed motion, to quantify motion on a single probe, with shanks in cortical and thalamic structures. We chose a superficial and a deep target because it is possible that motion affects different structures differently and wanted to investigate how stable the use of silicone adhesive is at reducing motion of tissue close to the craniotomy. Relative motion, extracted from voltage signals at cortical and thalamic sites, was minimal (Fig. 2a – middle). Motion was less than the size of a single electrode (<5 um) and close to the distance between sites (Fig. 2b).

**Figure 2.**
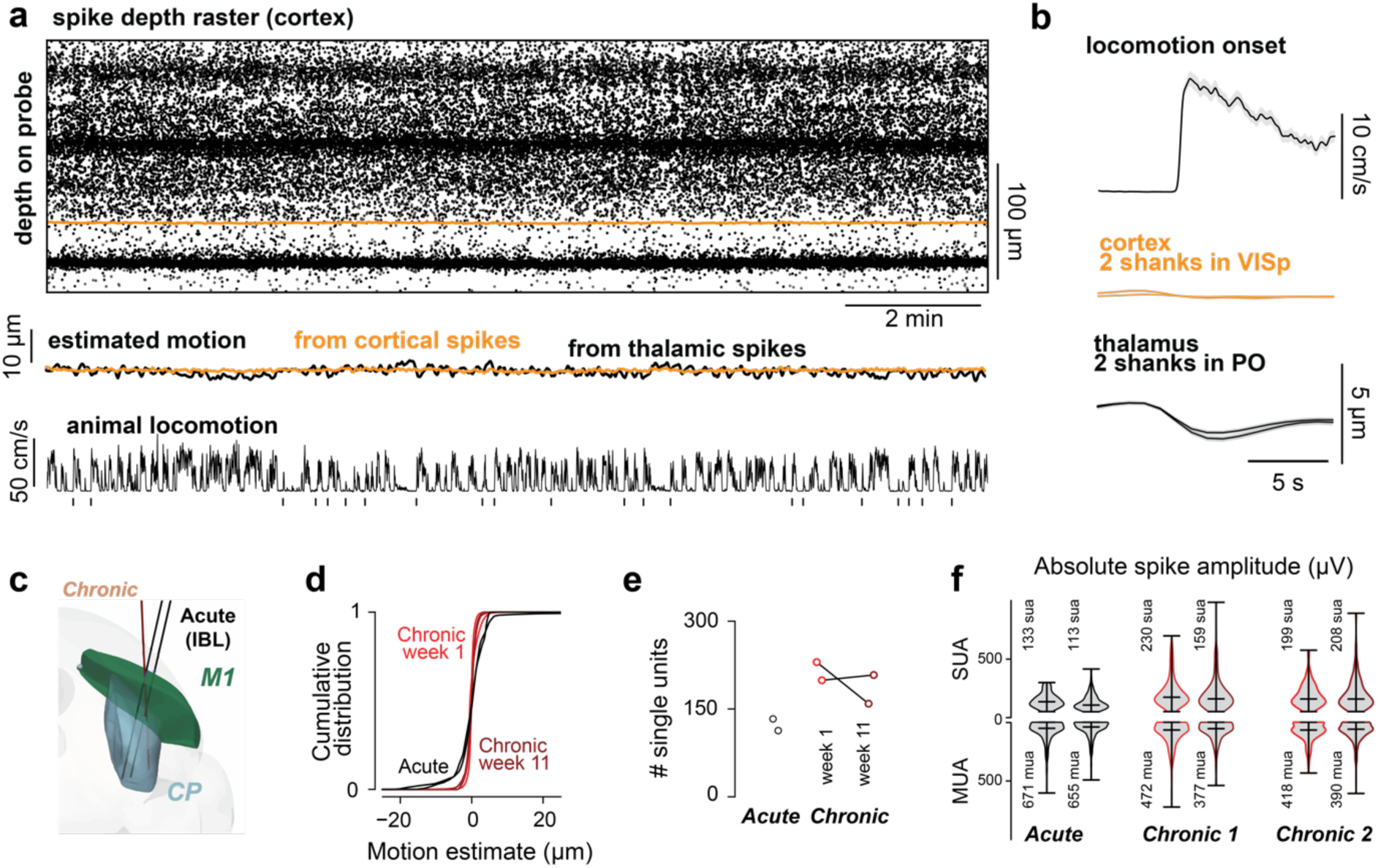
Minimal motion of the brain in relation to the shank and unit yield is comparable to published data. a) Top: Raster plot for a shank with recording sites in cortex (color indicates spike amplitude). Recordings with 4 shank probe (NPa - mouse in Fig.1d) with 2 shanks in cortex and 2 shanks in thalamus (PO). Middle: Motion estimates from DREDge for individual shanks in cortex and thalamus (black). Bottom: locomotion speed of the mouse on a treadmill; vertical markers indicate locomotion bouts with pauses longer than 5 seconds. There is no motion of the probe in relation to the brain tissue while the mouse engages in head-restrained locomotion. b) Average motion aligned to the onset of locomotion bouts. Estimated motion is smaller than the distance between sites in Neuropixels 2.0 probes (12µm). Brain motion in head restrained condition is less than 5 µm on locomotion onset. c) Planned chronic trajectories for comparison with available published datasets from the International Brain Laboratory that were acquired in the acute condition with the same behavioral task. d) motion in chronic (red - week 1; dark red - week 11) is reduced in comparison to the acute datasets (black). e) Chronic single unit yield with the same processing and metrics is higher than for recordings obtained with fresh craniotomies in an acute setting both in the first week of training and after 11 weeks. f) spike amplitudes are maintained after 11 weeks.

In a separate set of animals implanted with Neuropixels 1.0, we compared motion within single sessions in our dataset to sessions recorded in the same brain areas (the primary motor cortex and striatum), during the same behavioral task, and using similar electrodes but collected in an acute setting (International Brain Laboratory et al., 2023b). This allowed us to compare early (first week of training) and late sessions (∼2.5 months after) with published data. We compared the magnitude of motion of the brain in relation to the probe (Fig. 2d), the single unit yield (Fig. 2e) and the amplitude of single and multi-units (Fig. 2f). Importantly, we used the same sorting algorithm, without motion correction, and the same criteria for single unit selection. The criteria were: less than 0.1 false positive spikes, estimated from refractory period violations using 1.5ms as refractory period and 0.2ms as censored time (Hill et al., 2011; Llobet et al., 2022); less than 0.1 missed spikes, estimated from the amplitudes of individual spikes; principal waveform amplitude larger than 50µV, spike duration of the principal waveform longer than 0.1ms; exhibit spikes in over 60% of the recording (presence ratio). These criteria are slightly more stringent than the inclusion criteria specified in (International Brain Laboratory et al., 2023a). We selected these criteria to ensure that we obtained the same number of units when sorting with Kilosort 2.5 and 4.0 (Pachitariu et al., 2023) (Supplementary Fig. 2a). All data presented, including data from published work, was preprocessed using our custom pipeline and the same versions of open-source software packages. We provide containerized environments for reproducibility that include software for preprocessing, spike-sorting and all unit metric calculations used here (see Code Availability).

Our chronic recordings had less tissue movement in relation to the probe than IBL acute recordings (Fig. 2d). This was statistically significant in comparison to both early and late chronic sessions (acute-early: p<1e-10, acute-late: p<1e-10, Fligner-Killeen test).

We then set out to compare single unit yield. A major concern with chronic recordings is tissue health: unhealthy craniotomies or inflammation may reduce single unit yield due to gliosis or other immune factors (Hermann and Capadona, 2018; Xiang et al., 2024). The single unit yield on our recordings was comparable if not higher than that of IBL recordings (Fig. 2e). Some of this difference is in part due to the presence ratio criteria (60%) that we adopted. Acute sessions had 78±4 passing units more when the presence ratio was removed from the criteria whereas chronic sessions had 50±3 more single units (average ± s.e.m.). Despite the yield, the improved stability of our chronic device might come at the cost of worsened unit quality, e.g., a reduction in spike amplitudes due to inflammation. To test this, we compared spike amplitudes in our early and late chronic recordings with the IBL acute dataset. We found that spike amplitudes of single units are greater than IBL recordings made in the acute condition (p < 1e-10, one way ANOVA. p < .01 for all possible pairs of individual acute versus chronic sessions with Tukey’s HSD).

These results suggest that recordings using our approach are stable within single sessions and achieve similar, if not higher, yield to recordings obtained using standardized methods and protocols developed across laboratories (International Brain Laboratory et al., 2023a, 2023b).

### Implant stability and unit yield across sessions

We were motivated to develop this approach in part by the need to track neural activity during long timescales, such as during learning of perceptual decision-making tasks. Learning usually occurs over weeks to months depending on the difficulty of the task, so we set out to quantify stability over long timescales. We first quantified the motion of the brain in relation to the probe, and then the single unit yield across months.

Using the same animal as in Fig. 1d, we concatenated sessions acquired across 260 days (5 minutes for each session) and estimated the motion across the depth of cortex and thalamus, for two shanks. We found that inter-session movement depended on the brain region. There was more movement across sessions in the cortex (Fig. 3b, mouse 1: shanks 1 and 2) than in the thalamus (shank 3 and 4; but see also Supplementary Fig. 3). The median shift between sessions was 4.9±7.6 µm for cortex and 1.6±8.2µm for thalamus (median ± std). Interestingly, movement was more prevalent for the first month, suggesting that it might be related to recovery from surgery. This magnitude of motion can be easily corrected with registration algorithms and is less than the motion observed within acute sessions, in the most extreme cases (Windolf et al., 2023).

**Figure 3.**
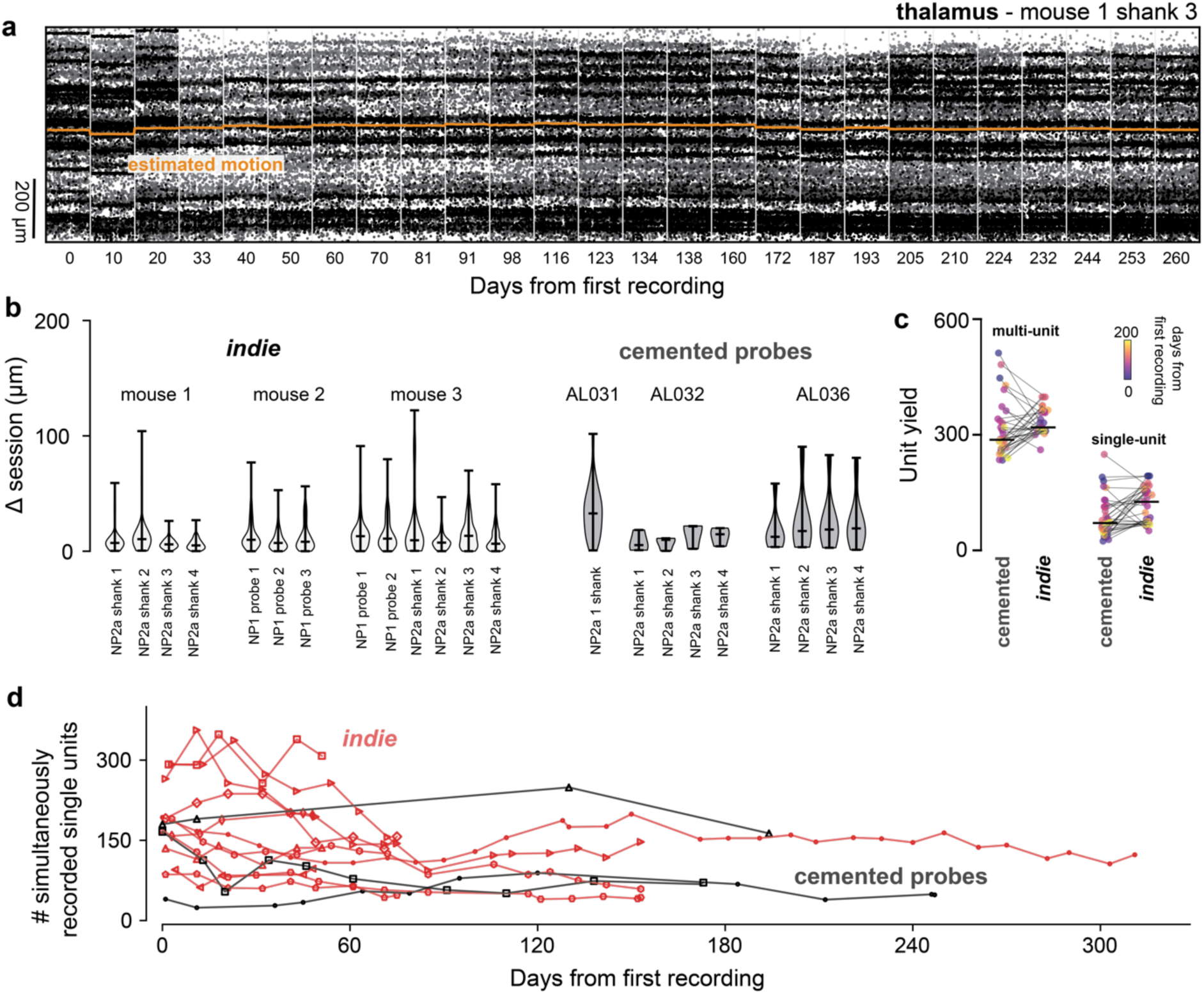
Motion across sessions and long-term stability is comparable to cemented probes. a) Spike raster and motion estimate across recording sessions for a single shank implanted in the thalamus (26 sessions are shown). Absolute estimated motion (orange) is overlaid on the raster. b) Median motion across sessions is comparable to that in cemented probes (Steinmetz et al. 2021). c) Multi unit and single unit yield per probe are comparable to cemented probes (matched to days from recording and only recordings passing through V1) N=3 mice for each group. d) Single unit yield on longitudinal experiments. Data in red are the single unit yield subsampled from Figure 4a. N = 5 mice for our fixture and N=3 for cemented probes.

The best possible scenario for tethered probe stability is when the probes are directly cemented to the skull. We then set out to compare the stability probes secured by our fixture with those of probes that were irreversibly attached to the skull (Lebedeva et al., 2020). Absolute probe motion between sessions was indistinguishable from motion in cemented probes (Fig. 3b - ANOVA p=0.178). We note that we had many more recording days with our fixture than sessions available in the cemented probe dataset and provide a comparison of absolute motion for roughly matched data points in Supplementary Fig. 3.

The electrodes in chronic recordings can remain implanted for months and therefore can evoke inflammatory responses that in turn encapsulate the electrodes and impact single unit yield. This can be an issue if the implant is not stable, leading to a decay of single unit yield. For cemented probes, this can be less than a log unit over the time course of months (Fig 2E in (Steinmetz et al., 2021)).

To measure the impact of our implant and optimized surgical procedures on single unit yield, we quantified single unit yield across months. Note that we use the “single unit” term with caution here, we selected units based on a fixed criteria as opposed to manually inspecting units because of the large number of recorded sessions. We classified “single units” using the criteria described above and compared yield using our implant with yield in the cemented probes dataset (Fig. 3c). We sub-sampled our recordings to match the interval between sessions available for cemented probes and included only insertions through the primary visual cortex, thus matching the regions recorded with the cemented probes. Reassuringly, recordings with our fixture had similar single unit yield to the dataset with cemented probes (119±8 for our fixture and 87±9 for cemented probes; mean±s.e.m., not significant KS p-value 0.11 - Fig.3c). Experiment variability may reflect experimenter, subject or procedural differences but the fact that our recordings can achieve high unit yield for sustained periods suggests that implants using the fixture are as stable as cementing the probe (Fig. 3d, but see also Fig. 4a).

**Figure 4.**
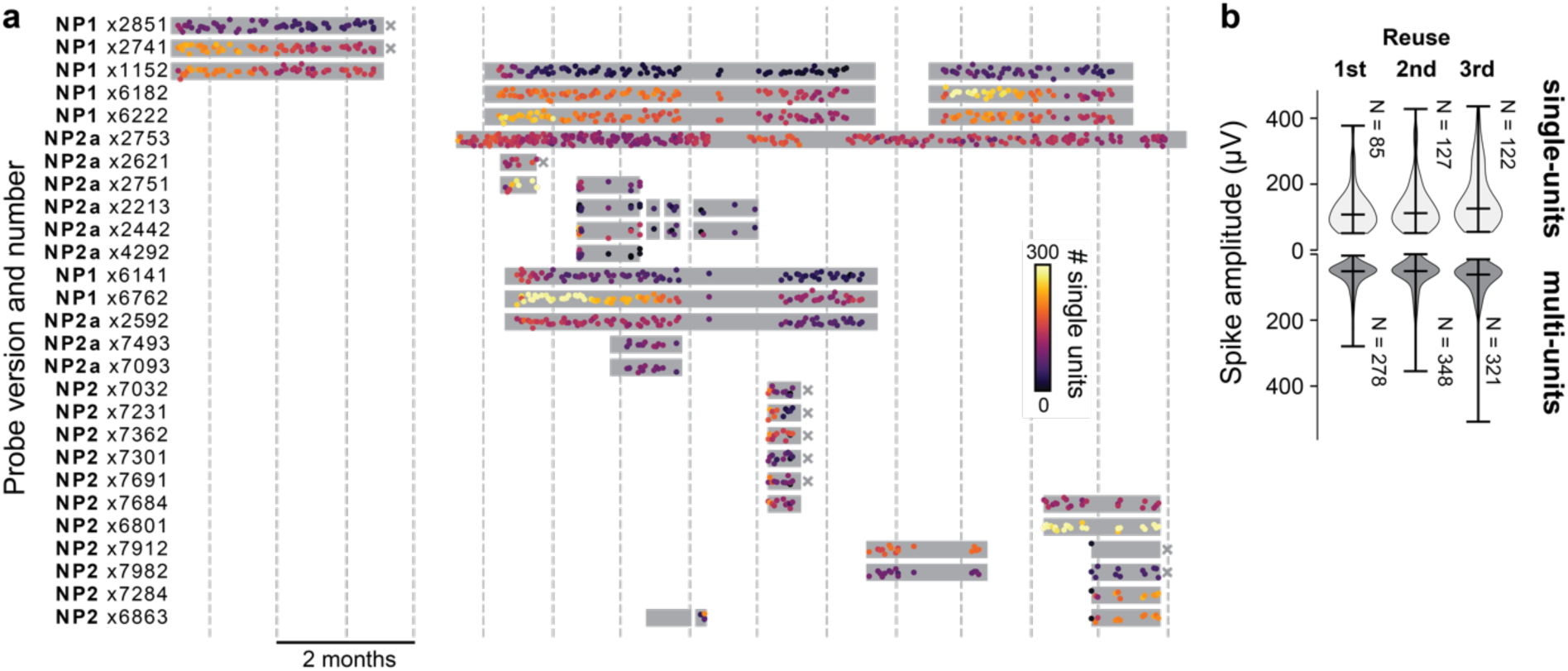
Probes can be reused without impairing the ability to detect single units. a) Experimental lifecycle for each of 27 probes. Gray boxes indicate the duration of an implant; each dot reflects a single recording session; colors indicate the number of units recorded during that session. “x” indicates failed extractions; all others are successful (NP2a x2753 is still implanted). Gray vertical lines: months. b) Average spike amplitude of the electrode with highest spike for multi units and single units recorded from the same areas on the 1st through 3rd reuse (2nd to 4th probe use). Number of single units is similar across reuses. Data are from NP2a x2442 and NP2a x2213.

### Probe extraction and reusability

A key requirement for scalable multi-probe recordings to be accessible to multiple laboratories is the re-usability of the probes. We therefore devised a mechanism to release the probe after an experiment and retrieve it unharmed. Importantly, our approach allows probes to be reused in acute experiments as well as re-implanted chronically because it does not permanently attach the probe to the fixture. Further, when multiple probes are implanted chronically, they remain physically separated (i.e., they are not glued together), affording the opportunity to re-use the probes in altogether different configurations in future experiments targeting new areas. To recover probes, the fixture screw is removed, and the probe retracted through the dovetail socket. This mechanism allowed us to recover most of the implanted probes.

We report on 42 chronic insertions, using 27 probes into 16 mice (2 mice with 1 probe, 7 with 2 probes, 4 with 3 probes, 2 with 4 probes and 1 with 6 probes) from which we collected 717 sessions and spike sorted 1381 recordings. All insertions were successful however one probe in a 4 probe implant failed 1 day after implantation. We recovered probes 32 times (4 mice had at least one unsuccessful recovery) and re-implanted 11 of those probes 16 times – all recovered probes were tested after recovery and were functional. All insertions and recording sessions are summarized in Fig. 4 (NP2a x2753 is still being recorded and is shown in Fig. 1d). One of the insertions of NP2 x6863 was in the same mouse/craniotomy and is shown in Fig. 5d.

**Figure 5.**
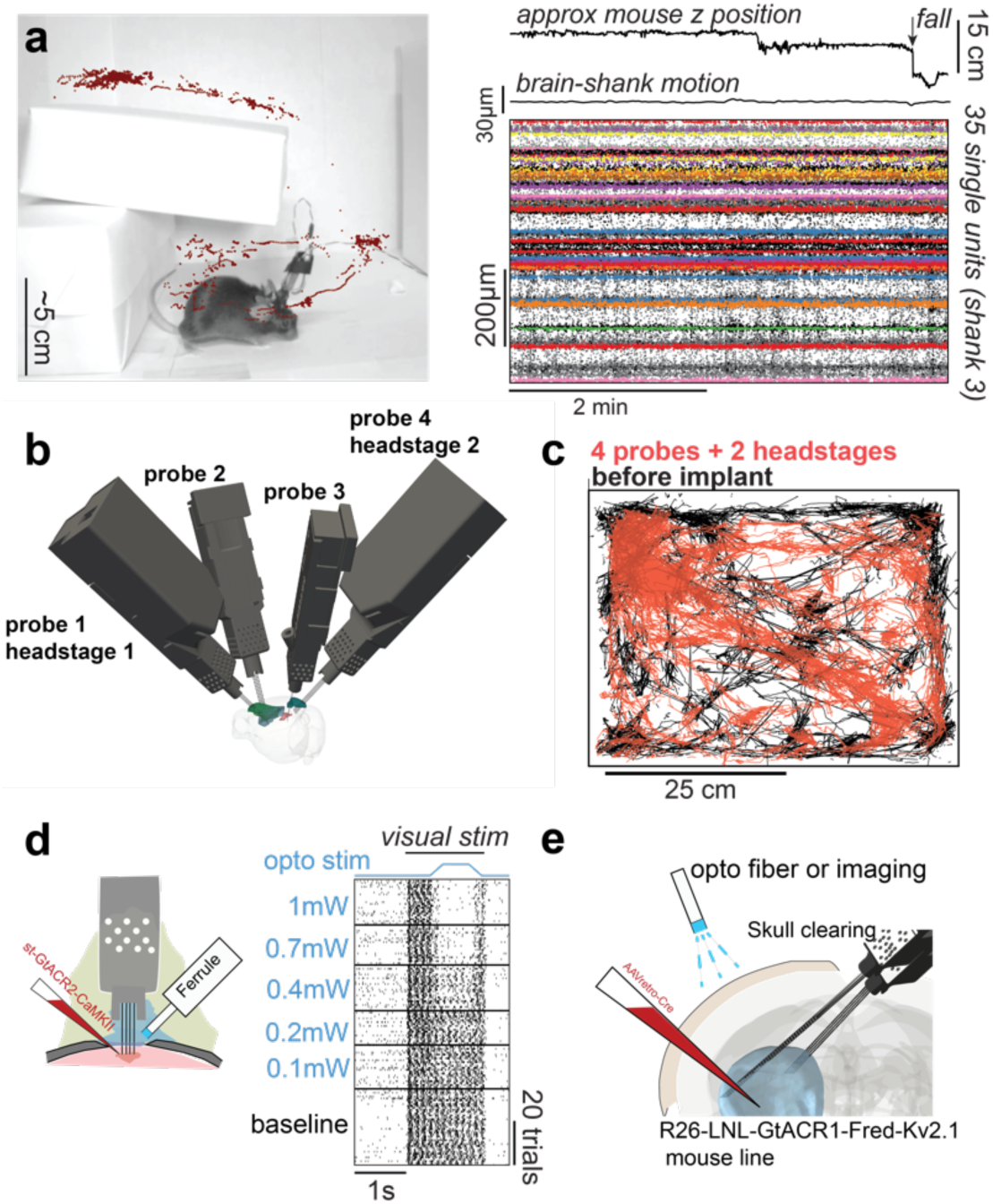
Stability during freely moving experiments, freely moving behavior with 4 probes and integration with optogenetics. a) Implants can be used in freely moving contexts. Recordings remain stable during natural exploration and a vertical drop of 15cm. top: approximate mouse z position in the environment and moment when the mouse takes a fall; middle: estimate of brain motion in relation to the shank. bottom: single unit spikes (color) overlayed on spike detection (black). b) Implant trajectories for a mouse with 4 probes and 2 headstages (∼3.5g without dental cement). c) Movement in an open arena during 45min. Black: mouse movement before implant. Red: Mouse movement with 4 probes and 2 headstages implanted. d) Implant flexibility enables recording and stimulating neurons with optogenetics from the same craniotomy. e) The implant also allows experimental approaches that were previously restricted to acute settings such as inserting probes chronically from the opposing hemisphere and stimulating cortical neurons through the intact cleared skull.

Importantly, these experiments were done during the development stage of the fixture and experimental procedures. This allowed us to optimize the protocols for implantation and extraction. A major change resulting from this optimization was on the selection of resin (initially we were using Black v4 which is more brittle and resulted in failure). Another identified failure mode was using too much thread-glue, which resulted in 3 probes being glued to the fixture. We recovered 2 probes with all shanks intact after drilling but these probes were not functional - these were the only cases where we used the drill during probe recovery. Notably, we did not recover 5 of the 6 NP2 probes implanted in the same mouse (at least in part) due to a defective stereotaxic holder which caused the probes not to be properly aligned to the axis of insertion (this is now corrected). We provide instructions to mitigate these issues in the methods section and are currently investigating ways of maximizing explant reliability. Nonetheless, we are confident that our implantation method is a major improvement over methods that rely on manipulators/stereotaxic equipment.

To assess the impact of probe reuse on our ability to isolate single units, we extracted 2 probes used in another experiment (implanted for 3 weeks, not shown because the implant targeted different brain areas) and re-inserted them consistently with the same trajectories in 3 different animals. We waited at least 10 days before each explant. Insertions targeted the same brain areas in the 2 hemispheres; data from the two hemispheres was pooled. Upon retraction, probes were placed in warm tergazyme for at least 6h, DS-2025 for at least 2h and deionized water for 1-2h. First, we looked at the Median Absolute Deviation (MAD) for individual channels. If the recordings contain more noise across re-insertions we expect MAD to increase. We did not observe an increase in MAD across re-insertions (11.43+-0.08 mean±s.e.m.). Next, we looked at single and multi-unit yield across re-insertions of the same probes. Single and multi-unit yield were not qualitatively distinguishable across re-insertions (Fig. 4b). Lastly, signal amplitude was not impacted by probe re-use because the spike amplitudes of single units were similar across insertions (Fig. 4b). In an additional set of experiments, we recovered 6 probes after being inserted for ∼6 months and 3 of those probes were reimplanted (NP1 x1152, NP1 x6182 and NP1 x6222) for an additional 3 months without visible decay in unit yield across re-insertions (see Fig. 4a).

Our results suggest that probes can be extracted and reused without impairing single unit yield using our fixture. Importantly, we were able to recover and reuse probes in longitudinal experiments (after ∼6 months of use) and our method of using the dovetail socket to recover probes does not rely on alignment with stereotaxic devices or manipulators and, therefore, allows recovering probes inserted at different angles in minutes.

### Use in freely moving contexts

One advantage of chronic implants over acute recordings is that they can be used in freely moving animals. The reduction in weight provided by our design not only allows more probes to be inserted simultaneously, but also invites use cases in freely moving contexts. We set out to test the severity of motion artifacts when mice were freely exploring an environment. We prepared an arena (Fig. 5a) that mice can explore in 3 dimensions. We wanted to test if there are artifacts when mice take a fall since this is one of the most severe tests of recording stability.

We incentivized a mouse to explore an environment using treats and tracked the animal’s position while recording neural activity. The z position (not corrected for perspective; Fig.5a) was extracted using pose estimation software (Mathis et al., 2018). We then quantified motion of the brain in relation to the electrodes and saw minimal motion during freely moving exploration (Fig.5a, ‘brain-shank motion’). We observed no artifacts due to animal motion in single unit activity (Fig.5a, right). Importantly, this was true even when the mouse took a fall of 15cm (Fig. 5a, arrow) demonstrating that the approach can be used to track cells in highly dynamic environments and even during falling.

For recording in freely moving conditions, each probe is connected to a headstage that also needs to be carried by the animal. This means that, for Neuropixels 2.0, mice will carry an additional ∼0.5g that enables recording from 2 probes. To facilitate freely moving experiments, we engineered a casing for the headstage that connects to the fixture. The headstage casing allows connecting to 2 probes implanted at independent angles (unlike Neuropixels 1.0, two 2.0 probes can share the same headstage). Previous freely moving experiments in mice have been limited to 2 probes, using our device we were able to record from 4 probes in freely moving settings (Fig. 5b). The combined weight of 4 Neuropixels 2.0 probes and 2 headstages, and the chronic holder assemblies is under 3.5g. To investigate whether the implant dramatically impaired the ability of mice to explore an environment, we tracked the position of a mouse in an open arena of 50 by 50cm using pose estimation software (Mathis et al., 2018) (Fig. 5c black).

We then implanted 4 probes in the left hemisphere of that mouse and connected them to 2 headstages that remained implanted. The tracked animal position while recording neural activity are reported for 13 days after surgery and with both headstages connected through standard twisted pair wires and without commutator. The mouse explored the entire arena to a similar extent as before the implant (Fig. 5c - red). We highlight that we purposefully positioned one of the headstages in the most anterior probe which is the configuration that imposes the greatest strain on the animal because the center of mass is more anterior. Our methods, therefore, enable freely moving experiments with 4 Neuropixels probes while minimally impacting naturalistic exploratory behaviors.

## Discussion

We developed a method to implant and recover multiple probes chronically in targeted locations of the mouse brain using a novel device, the *indie*. We report 6 probe simultaneous recordings (24 shanks) from the same hemisphere of the mouse brain in head-restrained contexts and 4 probes (16 shanks) in freely moving contexts. Importantly, we devised a strategy to independently set probe angles. This allows more probes to be inserted and enables experiments previously out of reach (in chronic settings) such as combining with optogenetics in the same craniotomy (Fig. 5d) or inserting probes chronically from the opposing hemisphere and stimulating/recording from the cortex through the cleared skull using optical methods (Fig. 5e).

### Comparison to other solutions

Our fixture tripled the number of probes simultaneously implanted in comparison to the state-of-the-art work in mouse (van Daal et al., 2021; Jones, 2023; Bimbard et al., 2023; Horan et al., 2024) and doubled the capacity in freely moving settings. Moreover, our new method for encapsulating the probes reduced the implant time to ∼1 hour/probe, a ∼3-fold improvement relative to (4 hours) published work (van Daal et al., 2021). Importantly, we were able to reduce surgery times using only standard stereotaxic surgery equipment; future work will optimize stereotaxic surgery equipment to further reduce surgery time by simultaneously implanting all probes, rather than implanting probes sequentially. Published alternatives that implant probes independently either have been designed for larger rodents (Luo et al., 2020) and are therefore much heavier than our solution; or require multiple parts implanted in succession rendering them, to our knowledge, impractical for implanting more than 2 probes in mice (Jones, 2023).

Despite using an independent fixture for each probe, our implant (NP1: 1.2g, NP2: 0.57g – probe included) is lighter than existing designs (Luo et al., 2020; van Daal et al., 2021; Bimbard et al., 2023; Horan et al., 2024) and allows implanting multiple probes in freely moving animals, without severely impacting naturalistic behaviors (Figure 5a,c).

In addition inter-session motion of brain tissue in relation to the probe shank(s) was indistinguishable from that observed in probes cemented over similar time durations (Steinmetz et al., 2021) and so was the single unit yield suggesting that our implants are as stable as permanently attaching probes to the skull with dental cement.

Our use of the dovetail in this fixture facilitated implant assembly and contributed to enhanced the ease of explant considerably (probes can be removed by hand without the need for a stereotaxic frame or manipulator).

### Caveats

Although our approach allows feasible implantation of up to 6 Neuropixels probes chronically in head fixed animals, implantation of this number in freely moving animals could pose some challenges. Specifically, because each pair of probes has a bespoke cable, implantation of 6 probes would require 3 cables which could be cumbersome for freely moving animals. Possible solutions include using a commutator to manage the cables, and the development of a headstage that is lighter and can accommodate more than two probes.

### Future directions

We see two natural extensions of this work. First, an appealing future direction for our approach would be to modify it for use in larger animals, such as rats or marmosets. We have assisted in implanting and recovering probes from rats successfully and with no modifications to the design but with a shield to protect the implant from direct impact. While we used a plastic shield, implants in larger animals could benefit from shields machined in metal, allowing easy autoclaving in advance of surgery. Interspecies differences in recording quality have not yet been characterized. Second, for mice, additional modifications could further lighten the total weight of the implant. For example, the dovetail on the probes could be shortened to save weight, provided that the extraction is not impacted. Another way to reduce weight is to shorten the height of the fixture; the current height is set to allow comfortable extraction. However, it seems feasible to, instead, create an external aiding mechanism for extraction. This would allow the probe encasing to be shorter and therefore lighter; effectively reducing the momentum acting over the head of the animal. Finally, advances in CMOS technology can reduce the size and weight of individual probes and decrease the number of headstages needed for multiprobe recordings (headstages are the heaviest part of the full assembly for freely moving recordings).

Our approach rests on the miniaturization and probe integration developments done by others (Jun et al., 2017; Steinmetz et al., 2021), a field that is under active development. Developments in probe technology will further open novel avenues to record at scale. Our method offers a flexible approach to maximize accessibility, reuse and extend the range of applications enabled by current probe technology.

## Methods

### Fixture fabrication

Design files and instructions are available at https://github.com/spkware/chronic_holder. The fixtures were printed using a Form 3+ resin printer (Formlabs). We selected the GreyPro resin for manufacturing, due to its optimal density, rigidity, and impact resistance properties. When setting up a print, we first used Inventor (Autodesk) to export “.stl” files from the original “.ipt” CAD files. These files were then loaded into PreForm (Formlabs) where printing orientations and support structures were defined. Fixtures were printed vertically and with the dovetail side facing the mixer side of the printer (multiple orientations were tested during prototyping and this was the most reliable). At the conclusion of 3D printing, parts were removed from the printing plate and washed for 15-20 minutes in isopropanol using the Form Wash tank (Formlabs). After washing, parts were allowed to dry for at least 4 hours. Parts were then UV-cured for 15 minutes at 80 degrees Celsius using the Form Cure (Formlabs). After curing, the support structures were carefully removed with tweezers and iris scissors. The screw-hole of the probe chassis was then hand-tapped with an M1 thread. Multiple dovetail tolerance values were printed, in the same platform, until the correct value was found. That is, when the probe could slide until 3mm from the bottom of the fixture with ease and tweezers were required to drive the probe to the bottom.

### Probe assembly and preparation

After probe fixture fabrication, the implant assemblies were prepared for surgery. We first used a dummy probe (or a non-functional real probe) to test the dovetail slide mechanism. This is done because the dovetail requires high dimensional accuracy and there can be dimensional variability across printing/washing rounds. When the dummy probe was confirmed to be properly held by the dovetail, we then inserted a real probe. Once the probe was in place, we secured it with an M1 screw. Loctite was applied to the screw threads only for the final turns of the screw. We discourage dipping the screw in loctite before screwing because loctite can flow into the dovetail mechanism, making removal more difficult. It is important to not overtighten or under tighten the screw, 1/4 to 1/3 turn once the initial resistance of hitting the dovetail occurs is sufficient to hold the probe in place for > 300 days. Before inserting the probe into the chassis, we soldered a silver wire (bare 0.01’’, 782500, A-M Systems) to the ground and reference contacts. We tested different configurations - in our hands connecting the ground and reference of the probe to a ground screw (M1) touching the dura was preferred to using internal reference. When using multiple probes, silver wires from all probes were connected to one ground screw. The ground screw was covered with silver epoxy to ensure electrical connection with the silver wires.

### Probe sharpening

Neuropixels probes were sharpened with an EG-45 Microgrinder (Narishige) before being inserted in the holder. We only sharpened probes once (before their first use). Probes were held at a 25-degree angle. Importantly, the probe was placed in the holder so that the backside of the shank (the side of the shank without electrical contacts) was facing the grinder. Then, the disk speed was set to 7 rotations per second. The probe was then lowered onto the spinning disk until there was a noticeable bend in the shank. Probes were left to sharpen for 5 minutes before being raised and observed under a stereoscope.

### Trajectory planning and visualization

Surgical trajectories were planned and rendered with custom software (Volume Visualization and Stereotaxic Planning, available at https://github.com/spkware/vvasp). The software relies on the PyVista project (Sullivan and Kaszynski, 2019) for rendering and includes a GUI for interactive surgical planning with a variety of brain atlases, downloadable through BrainGlobe (Claudi et al., 2020). It also includes geometries for a variety of probes and chronic probe holders that we developed to ensure that multi-probe implants will not result in holder collisions.

### Implantation surgery

Animal experiments followed NIH guidelines and were approved by the Institutional Animal Care and Use Committee of the University of California Los Angeles. We report results from implants in 14 male and 2 female mice (20-32g at time of surgery of different genetic backgrounds; C57BL6, FezF-CreER, stGtACR1xFezF or B6129SF1).

For some mice, headbar and Neuropixels implantation were performed within the same surgery. For other mice, these procedures were performed as separate surgeries, separated by a few weeks or months during which the mice were trained on behavioral tasks. For headbar implantation, the dorsal surface of the skull was first cleared of skin and periosteum. A thin layer of cyanoacrylate (VetBond, World Precision Instruments) was applied to the edges of the skull and allowed to dry. The skull was then scored with a scalpel or dental drill to ensure optimal adhesion. After ensuring the skull was properly aligned within the stereotax, craniotomy locations were marked by making a small etch in the skull with a dental drill. A titanium headbar was then affixed to the back of the skull with a small amount of glue (Zap-a-gap). The headbar and skull were then covered with Metabond, taking care to avoid covering the marked craniotomy locations. After the Metabond dried, the craniotomies for the probes and grounding screw were drilled. Once exposed, the craniotomies were kept continuously moist with saline.

The implants were held using the 3D-printed stereotax holder and positioned using a motorized micro-manipulator (Neurostar) and/or a manual stereotaxic manipulator (Stoelting) (in the case of multi-probe implants, we positioned, inserted, and cemented two probes at a time). After positioning the shanks at the surface of the brain, avoiding blood vessels, probes were inserted at slow speed (∼5 µm/s). The dura was left intact (when probes had difficulty penetrating the dura, repeated light poking of the dura with the probe tip eventually allowed insertion). Once the desired depth was reached, the craniotomy was dried and sealed with Dowsil 3−4680 (Dow Corning, Midland, MI). After the Dowsil 3-4680 dried, the craniotomy and shanks were sealed with Kwil-Sil (World Precision Instruments), completely enclosing the shanks, craniotomy, and the sealant enclosure in the fixture (Fig. 1a). The fixture was then secured to the skull with Metabond (C&B); mixed in a cold dish with 2 scoops to 4 drops of liquid and 1 of catalyst. After the Metabond dried, the stereotaxic arm was removed. This process was repeated until all probes were implanted. If a grounding screw was used, we connected it to the silver wire with silver epoxy. After all probes were implanted, additional layers of black Orthojet (Lang Dental) were applied to fully cover the fixture cement interface (Fig 1a). The mouse was then removed from the stereotax, and the 3D printed caps were secured to the fixtures. After surgical recovery, mice were treated with meloxicam and enrofloxacin for three days, then acclimated to head-restraint if required.

### Head-restrained recordings

Mice were habituated to head-restraint over a period of several days, where mice were head-restrained for increasingly long durations. After habituation, mice were trained to perform auditory or visual decision-making tasks. Animal behavior was captured with 3 video cameras (Chamaeleon3, FLIR). During the performance of these tasks, Neuropixels data was simultaneously acquired using SpikeGLX (https://github.com/billkarsh/SpikeGLX).

### Freely-moving recordings

Mice were placed atop a platform and allowed to freely roam a playground or an open arena. The position of the animal was recorded with a camera (Chamaeleon3, FLIR). Animal posture was extracted using DeepLabCut (Mathis et al., 2018) and depth estimated without correcting for the camera perspective. The posture labels were used to infer the location of the mouse in the arena. Neuropixels data was simultaneously acquired using SpikeGLX.

### Spike-sorting

Data were first preprocessed with a 300-12000Hz 3 pole butterworth bandpass filter, followed by ADC phase shift correction and common median subtraction. Sessions were spike-sorted with Kilosort4.0 (Pachitariu et al., 2023) using default parameters. To compute motion estimates using DREDGe (Windolf et al., 2023), we performed spike detection and localization with SpikeInterface (Buccino et al., 2020). Data aggregation and management were performed with DataJoint (Yatsenko et al., 2015).

Sorting jobs were run on a local GPU node, the Hoffman2 Shared Cluster provided by UCLA Institute for Digital Research and Education’s Research Technology Group and on Amazon Web Services using apptainer. Sorting jobs were automatically generated and executed by custom Python packages (https://github.com/jcouto/labdata-tools and https://github.com/spkware/spks).

### Probe recovery

To perform probe recovery, the mouse was first anesthetized with isoflurane or ketamine. The 3D printed caps were removed from the fixture, exposing the Neuropixels probe. We then cut the grounding wire ∼1cm away from the PCB and freed the flex cable from the tabs that secured it. We then loosened the fastening screw and slowly retracted the probe by pulling gently on the flex cable. The dovetail socket ensures that the shanks stay parallel to their insertion trajectory and do not break. Once the shanks were fully outside the brain and could be visualized, we carefully removed the probe from the fixture. Detailed step-by-step instructions are provided on the GitHub repository.

### Probe washing

After probe recovery, probes were washed in warm (<37deg) Tergazyme for ∼6 hours, followed by fresh DowSil-DS2025 for 2-8 hours to remove any layer of KwikSil that remained attached to the shanks. Finally, the probes were soaked in distilled water for 1-2 hours. Shanks were inspected under a stereo microscope; the procedure was repeated if the shanks were not clean.

## Code Availability

Designs and up-to-date instructions to build the fixtures are in https://github.com/spkware/chronic_holder

Code for preprocessing, sorting and unit metrics available and maintained in https://github.com/spkware/spks

Code for trajectory planning and 3D visualization of brain atlases, probes, and other objects available at https://github.com/spkware/vvasp

Code to reproduce figures in this manuscript available at https://github.com/spkware/chronic_recording_manuscript

An apptainer container with the installed software to perform spike sorting and analysis is available at https://figshare.com/articles/software/_b_i_indie_i_b_-_container_for_spike_sorting_and_analysis/26026384

## Author contributions

MDM and JC conceptualized and designed the fixtures, designed the experiments, built experimental rigs, optimized the surgical procedures, wrote code for preprocessing and data management, analyzed data and made figures, implanted all animals; and provided training. MDM, AK, MV, MBR and JC trained animals on behavioral tasks and collected data. MDM, AKC and JC wrote the manuscript. AKC provided resources, supervision and funding acquisition.

## Acknowledgements

We thank all members of the Churchland Lab for helpful discussions and common resources. UCLA DLAM for animal husbandry and colony maintenance. Chaoqun Yin for feedback on his implants on rats. Laura DeNardo for advice and equipment for tissue clearing. Trishala Chari and Carlos Portera-Cailiau for broken/dummy probes that we used at the start of the project for testing early designs. Anna Lebedeva and colleagues for releasing the raw data associated with Steinmetz et al. 2021 on figshare. Federico Sangiuliano Jimka and Daniel Aharoni for advice and support with 3d printing at the start of the project. Anjali Sinha and Maria Geffen for successful testing with the Form 2 printer, feedback on training instructions and notes. Mingmin Zhang (Weizhe Hong) and Arash Bellafard (Peyman Golshani), for feedback on their implants using our fixture. This work used computational and storage services associated with the Hoffman2 Shared Cluster provided by UCLA Institute for Digital Research and Education’s Research Technology Group. MBR was supported by a fellowship from the A.P. Giannini Foundation (20235719). This work was supported by an NSF-NCS collaborative award (2219946) and by NIH U19NS123716 and R01EY022979.

## Supplementary Figures

**Supplementary Figure 1.**
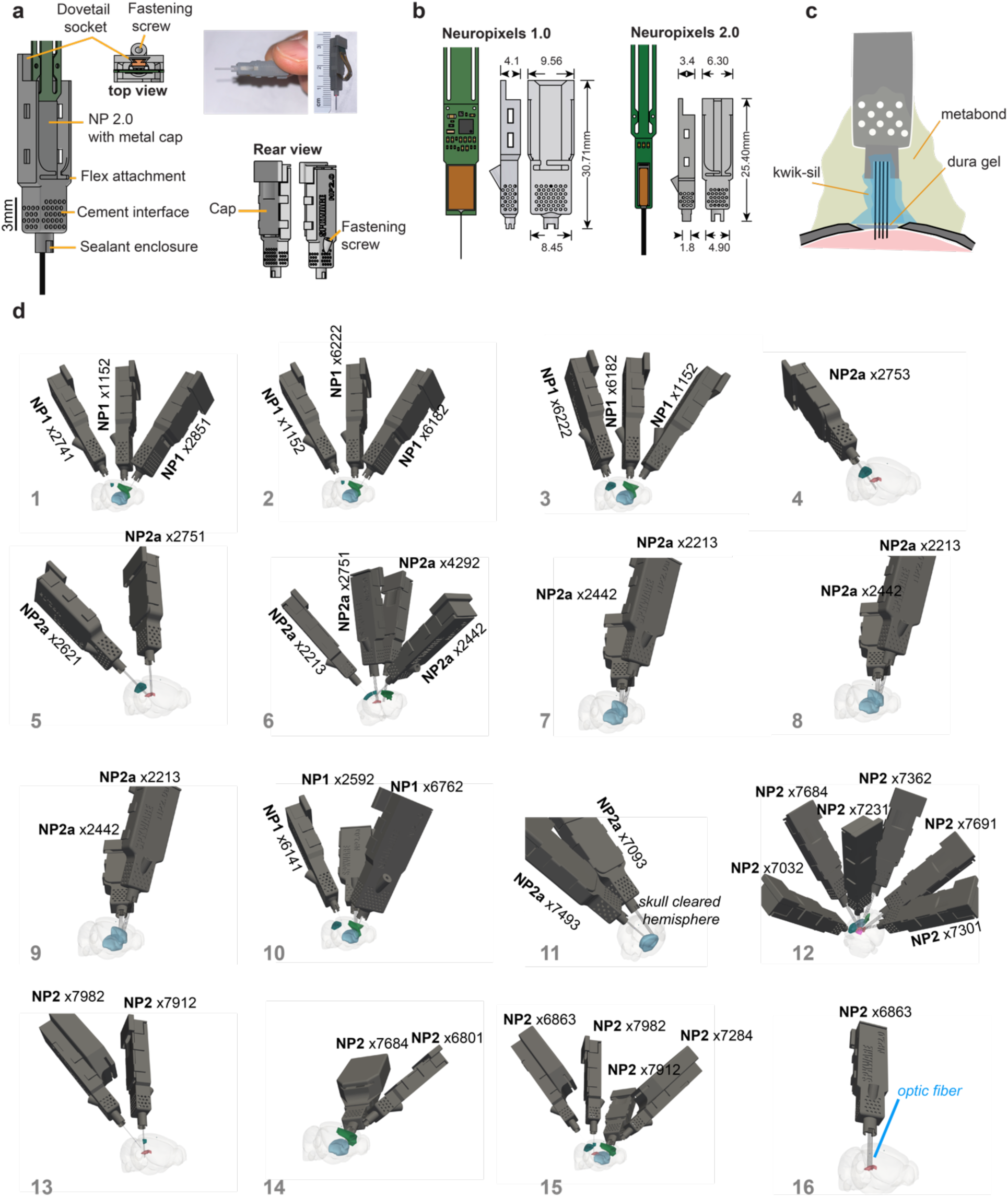
Schematics, surgery strategy and implants for all mice. a) Drawing of the probe fixture and assembly. b) Neuropixels 1.0 and 2.0 designs with dimensions. c) Surgical procedure schematic. d) Recording configurations for each mouse included in this study. The regions targeted for each experiment are also shown.

**Supplementary Figure 2.**
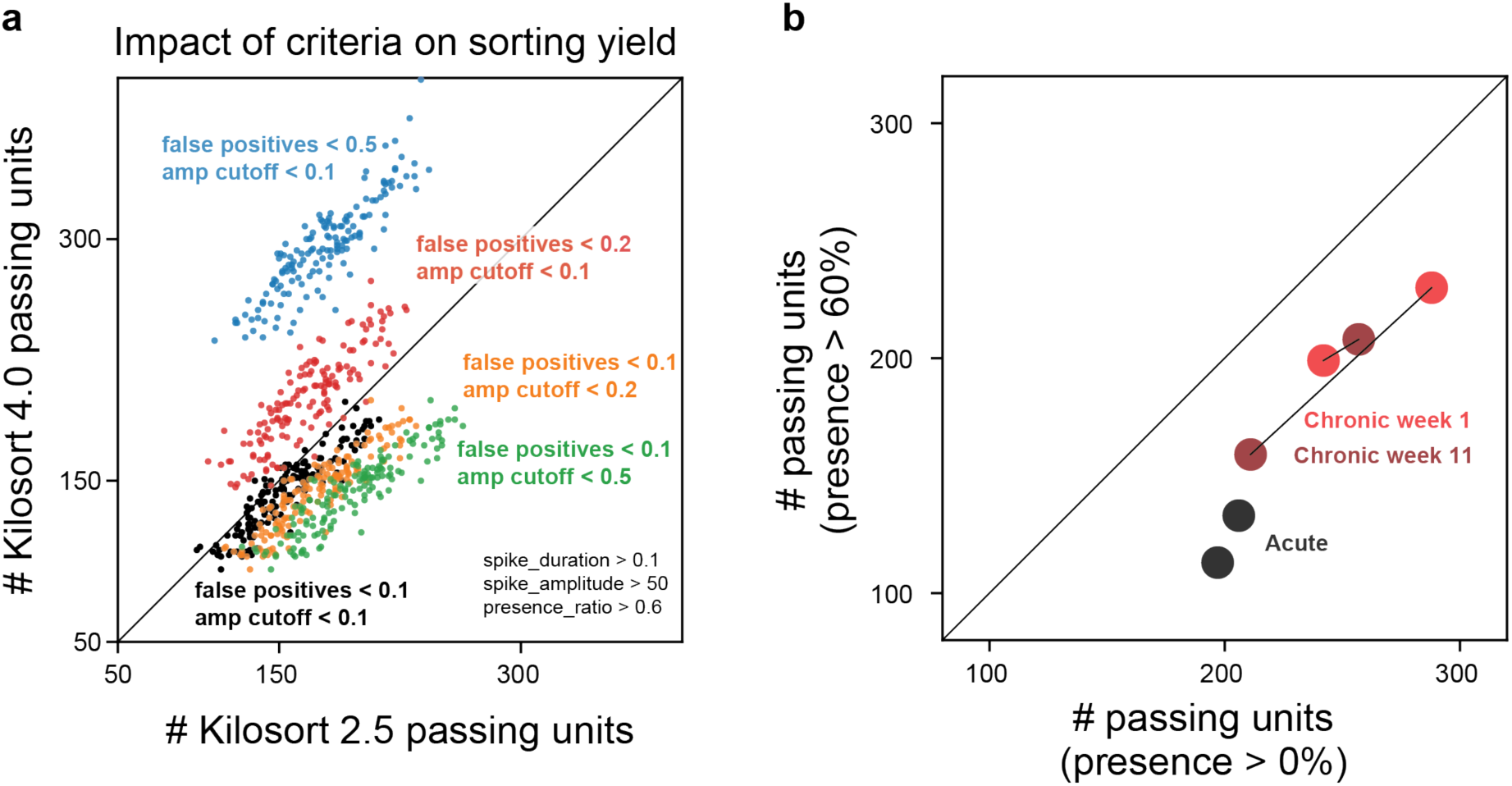
Impact of single unit criteria on sorting output depends on the kilosort version. To choose which kilosort version to use and single unit criteria to adopt, we plotted the number of passing units for kilosort 2.5 and kilosort 4.0.4. for all sessions in Figure 1d. Each color is the number of passing units with different criteria. We use only black (false positives < 0.1; amplitude cutoff < 0.1; spike duration > 0.1ms; spike amplitude > 50µV and presence ratio > 60%). Kilosort 4 output is insensitive to increases in amplitude cut off as reflected in the lateral shift from black, to orange, to green. This suggests that Kilosort 4 assigns more spikes to each unit. Interestingly, when we relax the criteria for false positives the number of Kilosort 2.5 units barely changes whereas the number of passing Kilosort 4 units increases, as seen in the upwards shift from black, to red, to blue. In this manuscript we used the most stringent criteria with Kilosort 4 output since it misses fewer spikes from each unit but these criteria capture slightly fewer units as Kilosort 2.5. b) IBL acute sessions have more units dropped than chronic sessions. This could be due to instability or relaxation of brain tissue around the shank during the recording session.

**Supplementary Figure 3.**
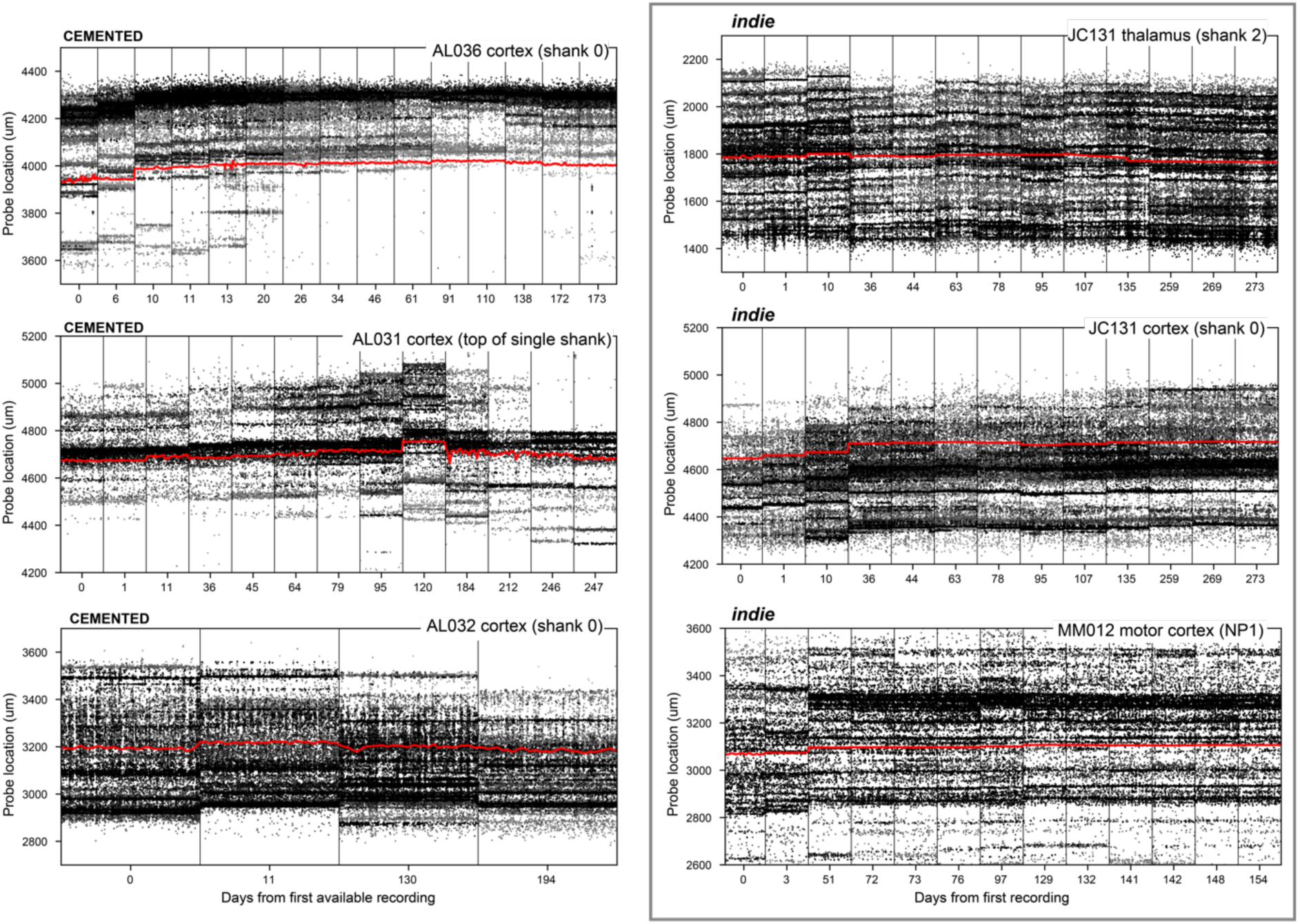
Motion of the brain in relation to the shanks is comparable to that of cemented probes. Red traces are quantified motion of the brain in relation to the shank over sessions. Left: recordings from cemented probes (Steinmetz et al 2021). Right: recordings with our fixture. We sampled our recordings to match the days for which there were data for AL031 as possible.

